# ATAD1 and the integrated stress response prevent clogging of TOM and damage caused by un-imported mitochondrial proteins

**DOI:** 10.1101/2023.09.06.556408

**Authors:** John Kim, Madeleine Goldstein, Lauren Zecchel, Hilla Weidberg

## Abstract

Mitochondria require the constant import of nuclear-encoded proteins for proper functioning. Impaired protein import not only depletes mitochondria of essential factors but also leads to toxic accumulation of un-imported proteins outside the organelle. Defects in mitochondrial protein import are associated with neurodegenerative and bioenergetic diseases. Here, we investigated the consequences of mitochondrial protein import stress in human cells. We demonstrated that un-imported proteins can clog the mitochondria by stalling inside the translocase of the outer membrane (TOM). We found that the integrated stress response (ISR) acted as the first line of defense to mitochondrial clogging by attenuating global protein translation and preventing excessive accumulation of un-imported proteins. A second mechanism, mediated by a mitochondrial ATPase, ATAD1, acted specifically to remove proteins from TOM and clear the entry gate into the mitochondria. ATAD1 interacted with both TOM and stalled proteins, and its knockout resulted in extensive accumulation of mitochondrial precursors as well as decreased protein import. Increased ATAD1 expression improved tolerance of cells to defective mitochondrial protein import, demonstrating the importance of this quality control pathway in surveilling protein import and its contribution to cellular health.

## Introduction

Mitochondria are essential organelles that are required for biosynthetic pathways of many metabolites including iron-sulfur clusters and ATP^1^. These organelles also play a central role in programmed cell death and cellular signaling^1^. The human mitochondrial proteome consists of approximately 1100 proteins, of which 99% are encoded by nuclear genes and synthesized on cytosolic ribosomes^2,3^. Hence, the constant import of proteins into the mitochondria is essential for maintaining the function of these organelles. The vast majority of mitochondrial proteins are first inserted into the translocase of the outer membrane (TOM) and are subsequently targeted to mitochondrial sub-compartments through various mechanisms^4^. In the case of matrix proteins, the route into the mitochondria is through TOM and then the translocase of the inner membrane (TIM). These proteins are translated as precursors with an N-terminal mitochondrial targeting sequence (MTS) that is cleaved by the mitochondrial processing peptidase (MPP) upon reaching the matrix. As protein import into the mitochondria often requires ATP and polarization of the inner mitochondrial membrane, it is susceptible to bioenergetic changes and various mitochondrial defects^4–6^. In addition, pathogenic mutations in protein import machinery genes, such as TIMM50 and acylglycerol kinase (AGK), directly affect delivery of proteins into the mitochondria^6–8^. When left unchecked, perturbations in protein import disrupt mitochondrial homeostasis by inhibiting the supply of newly synthesized proteins, as well as leading to toxic accumulation of un-imported proteins in the cytosol^9–11^.

Given the central role of mitochondria in the eukaryotic cell, various quality control pathways have evolved to cope with and repair dysfunctional mitochondria^12^. These include clearance of mitochondria or damaged components via mitophagy and mitochondrial-derived vesicles (MDVs), as well as activation of stress responses, such as the mitochondrial unfolded protein response (UPR^mt^)^12,13^. Additionally, the integrated stress response (ISR), which is known to be triggered by different stress stimuli, is induced by mitochondrial dysfunction^14–16^. The ISR attenuates global protein translation through phosphorylation of the eukaryotic translation initiation factor 2α (eIF2α) and induces transcriptional reprogramming by preferentially translating transcriptional regulators^15^. Recent studies have identified transcriptional stress responses that are triggered specifically by impaired mitochondrial protein import^9,11,17–23^. These pathways reduce translation and increase proteasomal degradation capacity to combat the accumulation of un-imported mitochondrial proteins in the cytosol.

Evidence from work in *S. cerevisiae* revealed that mitochondrial proteins can also accumulate inside the TOM channels when import is impaired, clogging the entry gate of the mitochondria^20,21,24–26^. Quality control pathways were demonstrated to alleviate this damage by extracting stalled proteins from TOM^21,24,25^. Our previous work discovered a role for Msp1, a mitochondrial outer membrane ATPase Associated with diverse cellular Activities (AAA), in clearing TOM channels, as part of the mitochondrial compromised protein import response (mitoCPR)^21^. While Msp1 is recruited to the TOM complex only under protein import stress, a different AAA, Cdc48/p97, and its adaptor Ubx2, constantly safeguard TOM against stalling^24^. The consequences of inefficient mitochondrial protein import are mostly unknown in mammals, yet recent studies demonstrated that TOM can be clogged by pathogenic mutants of mitochondrial proteins^26^. Stalling is not restricted to mitochondrial proteins, as amyloid precursor protein (APP) is mistargeted to mitochondria and clogs TOM in Alzheimer’s disease patients^27^. It is still unclear whether cellular pathways that repair mitochondrial protein import defects and clogging exist in mammalian cells. The mammalian Msp1 homolog, ATPase family AAA domain containing 1 (ATAD1), shares functional similarities to Msp1 in its ability to extract mistargeted proteins from the mitochondrial outer membrane^28–30^. However, a role for ATAD1 during mitochondrial protein import stress and its involvement in clearing TOM is yet to be discovered.

Here, we investigated the consequences of impaired mitochondrial protein import in human cells and how they are repaired. We demonstrated that impaired protein import can lead to accumulation of mitochondrial precursors in the cytosol and inside import channels. Two surveillance pathways were identified that help cope with damage caused by inefficient mitochondrial protein import. First, the ISR is activated to prevent further accumulation of un-imported mitochondrial precursors. Second, an ATAD1-mediated quality control mechanism directly acts to extract stalled precursors from mitochondrial import channels. We showed that the absence of ATAD1 leads to higher levels of un-imported proteins and exacerbates protein import defects. Finally, we demonstrated that both ISR and ATAD1 contribute to the overall fitness of cells with chronic mitochondrial dysfunction.

## Results

### Activation of the integrated stress response prevents accumulation of un-imported mitochondrial precursors

To investigate the fate of mitochondrial un-imported proteins under mitochondrial stress, we treated HEK293T cells with carbonyl cyanide m-chlorophenyl hydrazine (CCCP) and valinomycin^31^. These ionophores disturb the mitochondrial membrane potential and consequently block protein import. To assess mitochondrial protein import efficiency, we measured the import of the matrix protein ornithine transcarbamylase (OTC) in cells that stably express OTC-V5. The N-terminal mitochondrial targeting sequence (MTS) of OTC is cleaved upon import, allowing a separation of the un-imported precursor from the mature form of this protein by SDS-PAGE^4,32^. In untreated cells, the precursor of OTC-V5 was not detected due its efficient import and cleavage in the mitochondria (**Fig. 1A**). Surprisingly, inhibiting mitochondrial protein import by CCCP and valinomycin for 6 and even 24 hours resulted in a very mild accumulation of OTC-V5 precursor (**Fig. 1A**). Treatment with CCCP/valinomycin had a similar effect on the mitochondrial protein ATP5G1-HA (**Fig. 1B**). Notably, the precursor of transiently transfected ATP5G1-HA was visible even in untreated cells, which is consistent with previous reports^31^. This is presumably due to a combination of high expression levels and relatively low import efficiency. The short half-life of ATP5G1-HA in the mitochondria resulted in a complete loss of its mature form following 24 hours of CCCP/valinomycin treatment, demonstrating that mitochondrial protein import is indeed inhibited under these conditions (**Fig. 1B**).

**Figure 1.**
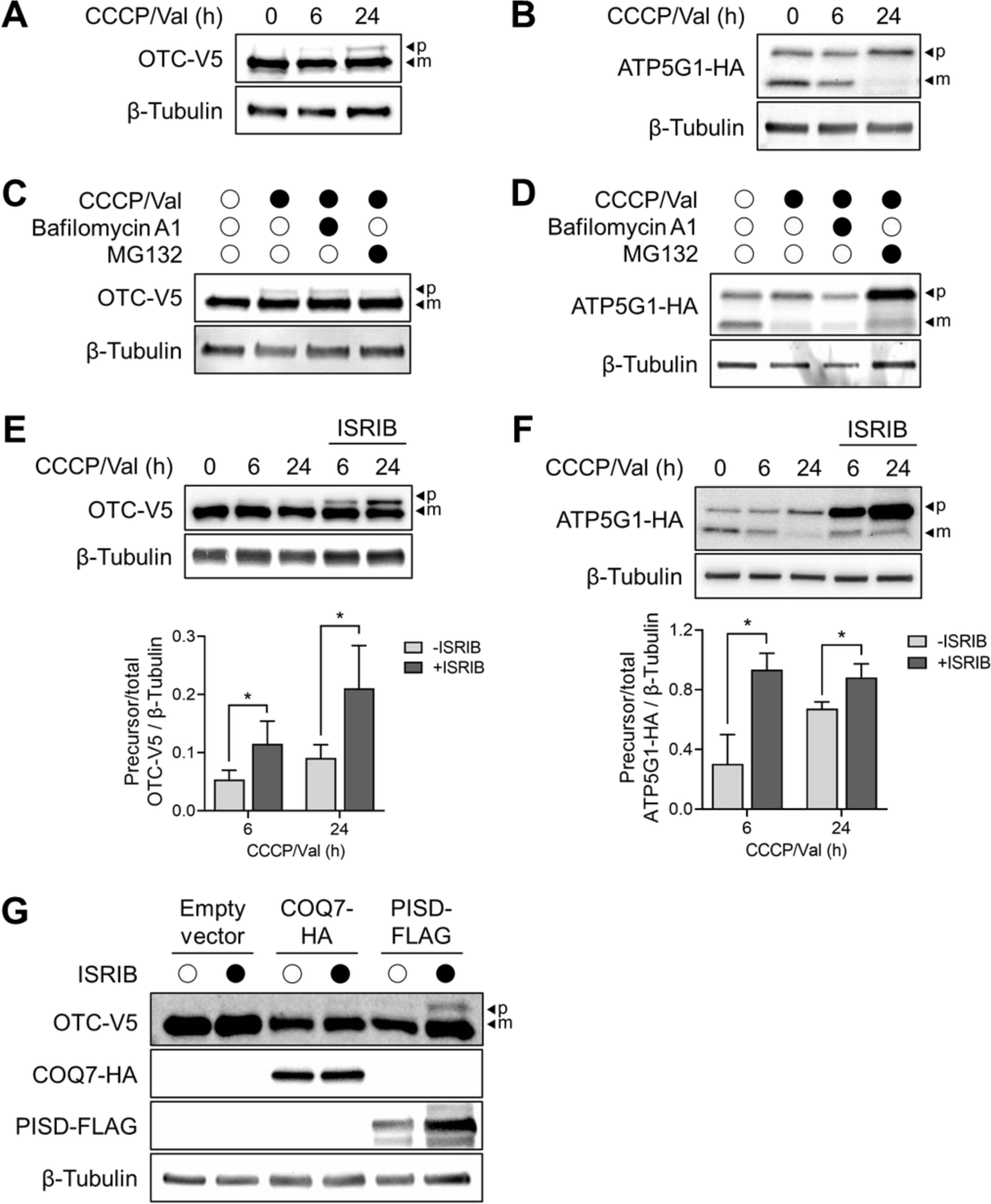
Un-imported precursors accumulate upon inhibition of mitochondrial protein import and the ISR. (**A**) Western blot analysis of OTC-V5 stably expressing HEK293T cells untreated or treated with CCCP (10 µM) and valinomycin (Val) (1 µM) for either 6 or 24 hours. (**B**) Western blot analysis of ATP5G1-HA transiently expressing HEK293T cells untreated or treated with CCCP (10 µM) and valinomycin (1 µM) for either 6 or 24 hours. (**C**) Same as A. Bafilomycin A1 (BafA1) (0.1 µM) and MG132 (20 µM) were added to the cells 6 hours prior to cell harvesting (CCCP/valinomycin treatment was for 24 hours). (**D**) Same as B. BafA1 (0.1 µM) and MG132 (20 µM) were added to the cells 6 hours prior to cell harvesting (CCCP/valinomycin treatment was for 24 hours). (**E**) OTC-V5 stably expressing HEK293T cells were either untreated or treated with CCCP and valinomycin in the absence or presence of ISRIB (0.5 µM) for 6 or 24 hours. Quantifications of OTC-V5 precursor levels normalized to total OTC-V5 and β-tubulin are shown (*n*=4). Data are mean ± SD. Paired t-test was used (* = *p* < 0.05). (**F**) ATP5G1-HA transiently expressing HEK293T cells were either untreated or treated with CCCP and valinomycin in the absence or presence of ISRIB (0.5 µM) for 6 or 24 hours. Quantifications of ATP5G1-HA precursor levels normalized to total ATP5G1-HA and β-tubulin are shown (*n*=3). Data are mean ± SD. Paired t-test was used (* = *p* < 0.05). (**G**) HEK293T cells were transiently transfected with OTC-V5 and either empty vector, COQ7-HA (control), or PISD-FLAG. *p*, precursor; *m*, mature protein.

What mechanisms prevent the accumulation of mitochondrial precursors under protein import stress? To address this question, we first tested whether protein degradation pathways act to eliminate un-imported mitochondrial proteins. The involvement of autophagy or mitophagy in mediating the clearance of mitochondrial precursors was excluded by treating cells with CCCP/valinomycin and the lysosomal degradation inhibitor bafilomycin A1 (BafA1) (**Figs. 1C and 1D**). Proteasomal degradation was previously demonstrated to act as a quality control mechanism for clearance of un-imported mitochondrial precursors^18,20,21,24,33^. To examine a possible role for the proteasome in eliminating mitochondrial precursors, we treated cells with CCCP/valinomycin and the proteasomal inhibitor MG132. No significant change in the accumulation of OTC-V5 precursor was observed following this treatment (**Fig. 1C**). The accumulation of ATP5G1-HA precursor was enhanced upon proteasomal inhibition when protein import was disturbed (**Fig. 1D**). This result is consistent with a previously reported role for ubiquilin chaperones and proteasomal degradation in quality control of ATP5G1 precursor in the cytosol^31^. Overall, our data suggest that mechanisms other than protein degradation pathways also exist to prevent accumulation of un-imported mitochondrial precursors.

We then asked whether the accumulation of mitochondrial precursors in CCCP/valinomycin-treated cells is controlled at the translational level. The ISR is induced by various mitochondrial stressors in mammalian cells, including CCCP^16,34–38^. ISR activation leads to enhanced phosphorylation of eIF2α and subsequent attenuation of global translation^15,35^. To examine the impact of ISR on precursor accumulation, we treated cells with CCCP/valinomycin and the ISR inhibitor (ISRIB). Inhibition of the ISR led to significantly increased levels of both OTC-V5 and ATP5G1-HA precursors following 6 and 24 hours of protein import stress (**Figs. 1E and 1F**).

CCCP and valinomycin cause severe mitochondrial damage and can affect other cellular membranes. We previously reported that in yeast, overexpression of mitochondrial proteins that contain a bipartite signal, such as Psd1, overloads the import machinery and leads to protein import stress^21^. We therefore used this non-pharmacological approach to validate our findings by co-expressing OTC-V5 together with the human Psd1 homolog, PISD, or a control mitochondrial protein, COQ7. Overexpression of PISD, but not COQ7, led to accumulation of OTC-V5 precursor in the presence of ISRIB, supporting that excess bipartite proteins impair mitochondrial protein import and activate the ISR (**Fig. 1G**). Consistently, phosphorylation of eIF2α was elevated in PISD-overexpressing cells (**Fig. S1**). Hence, our data suggest that ISR-mediated translation attenuation helps reduce the burden caused by accumulation of un-imported proteins in both pharmacological- and genetic-induced mitochondrial import stress.

### Mitochondrial precursors stall inside TOM under protein import stress

Next, we asked where un-imported mitochondrial proteins accumulate in the cell. We detected both mature and precursor forms of OTC-V5 in a mitochondria-enriched fraction of cells induced with protein import stress (CCCP/valinomycin/ISRIB), while almost no protein was detected in the cytosolic fraction (**Fig. 2A**). Association of precursors with mitochondria can indicate defects in either import into the organelle or MTS cleavage in the matrix. To distinguish between these possibilities, we treated mitochondria from stressed, OTC-V5 expressing cells, with proteinase K. The precursor, but not mature form, of OTC-V5 exhibited proteinase K sensitivity (**Fig. 2B**). Since OTC was detected using a C-terminal V5 tag, we conclude that at least the C-terminus of OTC precursor resides at the mitochondrial surface facing the cytosol. Similarly, both ATP5G1-HA precursor and mature forms were detected in the mitochondrial fraction, where only the precursor was accessible to proteinase K (**Fig. S2**). Interestingly, a population of ATP5G1 precursor was also detected in the cytosolic fraction (**Fig. S2**). These results are in agreement with previous studies, suggesting that the fate of un-imported mitochondrial precursors can differ between proteins^19,33,35^.

**Figure 2.**
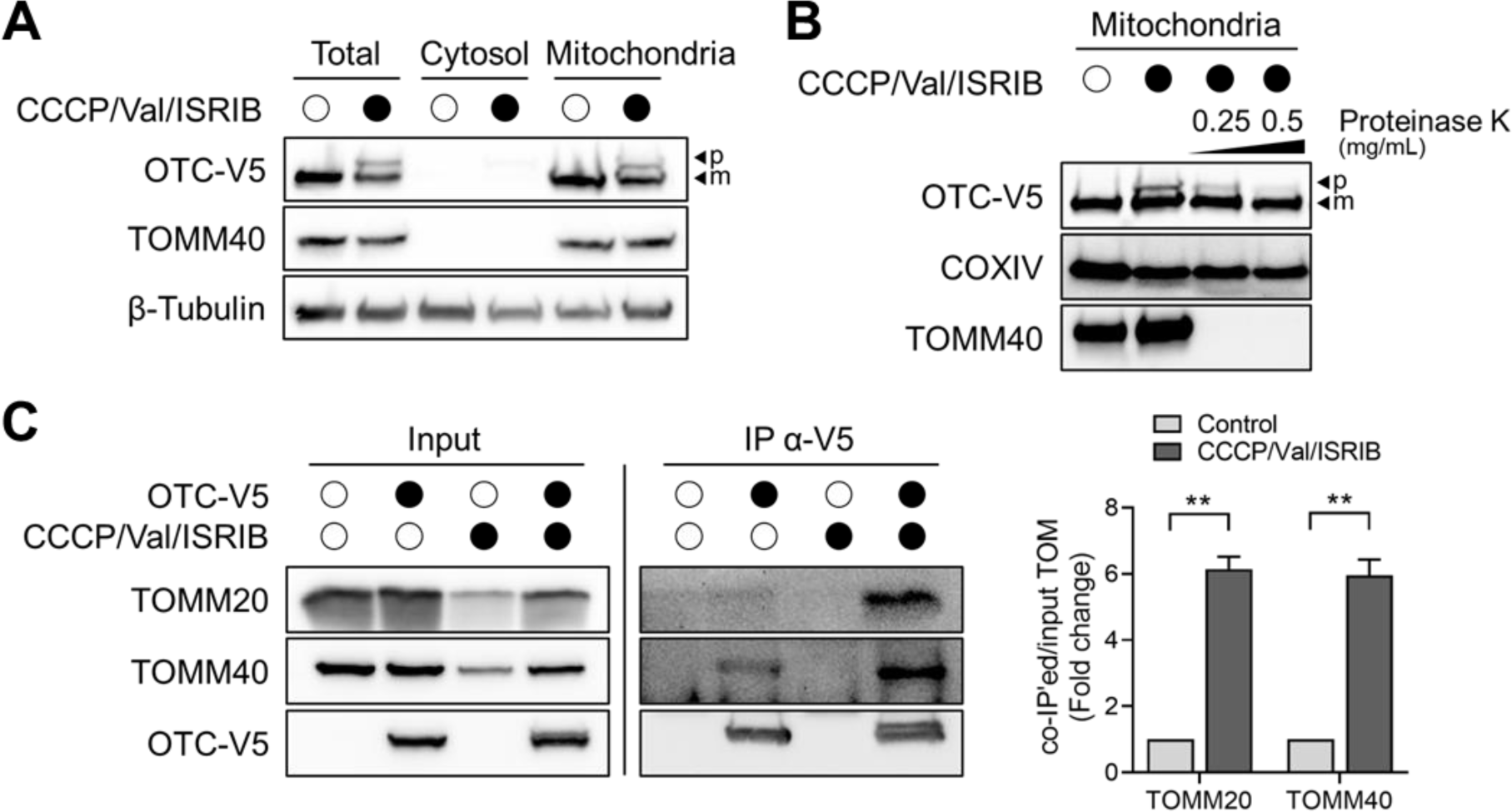
Mitochondrial precursors are stalled in TOM when mitochondrial protein import is impaired. (**A**) Mitochondrial and cytosolic fractions were enriched by differential centrifugation from OTC-V5 stably expressing HEK293T cells untreated or treated with CCCP (10 µM), valinomycin (Val) (1 µM), and ISRIB (0.5 µM) for 24 hours to induce protein import stress. TOMM40 was used as a mitochondrial marker. (**B**) Mitochondrial fractions from A were treated with 0.25 mg/mL or 0.5 mg/mL of proteinase K. TOMM40 was used as a mitochondrial outer membrane marker. COXIV was used as a mitochondrial matrix marker. (**C**) OTC-V5 was immunoprecipitated (IP) from HEK293T cells stably expressing OTC-V5, treated as in A, using antibodies to V5. Quantifications of co-immunoprecipitated TOMM20 and TOMM40 normalized to their respective inputs are presented as fold change, for which the control condition was set to 1 (*n*=3). Data are mean ± SD. Paired t-test was used (** = *p* < 0.01). *p*, precursor; *m*, mature protein.

Having established that the precursor of OTC accumulates at the mitochondrial surface when protein import is impaired, we next asked whether it is stalled inside TOM. Immunoprecipitation (IP) of OTC-V5 from untreated cells revealed a weak association of OTC-V5 with two components of the TOM complex: the import receptor, TOMM20, and the channel-forming subunit, TOMM40 (**Fig. 2C**). This association presumably represents the transient interaction of OTC with the import machinery while it is translocated into the mitochondria. Notably, upon induction of protein import stress (CCCP/valinomycin/ISRIB), the interaction between OTC-V5 and the TOM components was stabilized. This result suggests that when efficient entry into the mitochondria is prevented, precursors can clog the import channels in human cells.

### The ATPase ATAD1 prevents the accumulation of mitochondrial precursors during protein import stress

Do mechanisms other than ISR exist to prevent accumulation of precursors and clogging of mitochondria? Our previous work in yeast has shown that during protein import stress, Msp1 is recruited to TOM to mediate the extraction of stalled proteins^21^. We thus asked whether the human Msp1 homolog, ATAD1, plays a similar role in mitochondrial precursor quality control. To this end, we used CRISPR/Cas9 to delete *ATAD1* in HEK293T cells and observed an increase in the accumulation of OTC-V5 precursor following induction of protein import stress by CCCP/valinomycin/ISRIB treatment (**Fig. 3A**). The impact of *ATAD1* deletion on precursor accumulation was detected as early as 8 hours following induction of protein import stress (**Fig. 3B**). In addition to the elevated levels of OTC-V5 precursor, *ATAD1* knockout resulted in a reduced level of mature OTC-V5, suggesting that ATAD1 helps maintain a certain level of protein import (**Fig. 3A**). Notably, accumulation of mitochondrial precursors was not detected in *ATAD1* knockout cells under non-stress conditions, excluding the possibility that these cells have a general defect in protein import. The defects leading to OTC precursor accumulation in the knockout cells were restored by ectopic expression of wild type ATAD1, but not by the “substrate-trap” mutant, ATAD1^E193Q^ (**Fig. 3C**). ATAD1^E193Q^ contains a point mutation in the Walker B motif of the AAA domain and is therefore unable to hydrolyze ATP^28^. Thus, ATAD1 ATPase activity is essential for limiting the accumulation of mitochondrial precursors when protein import is impaired.

**Figure 3.**
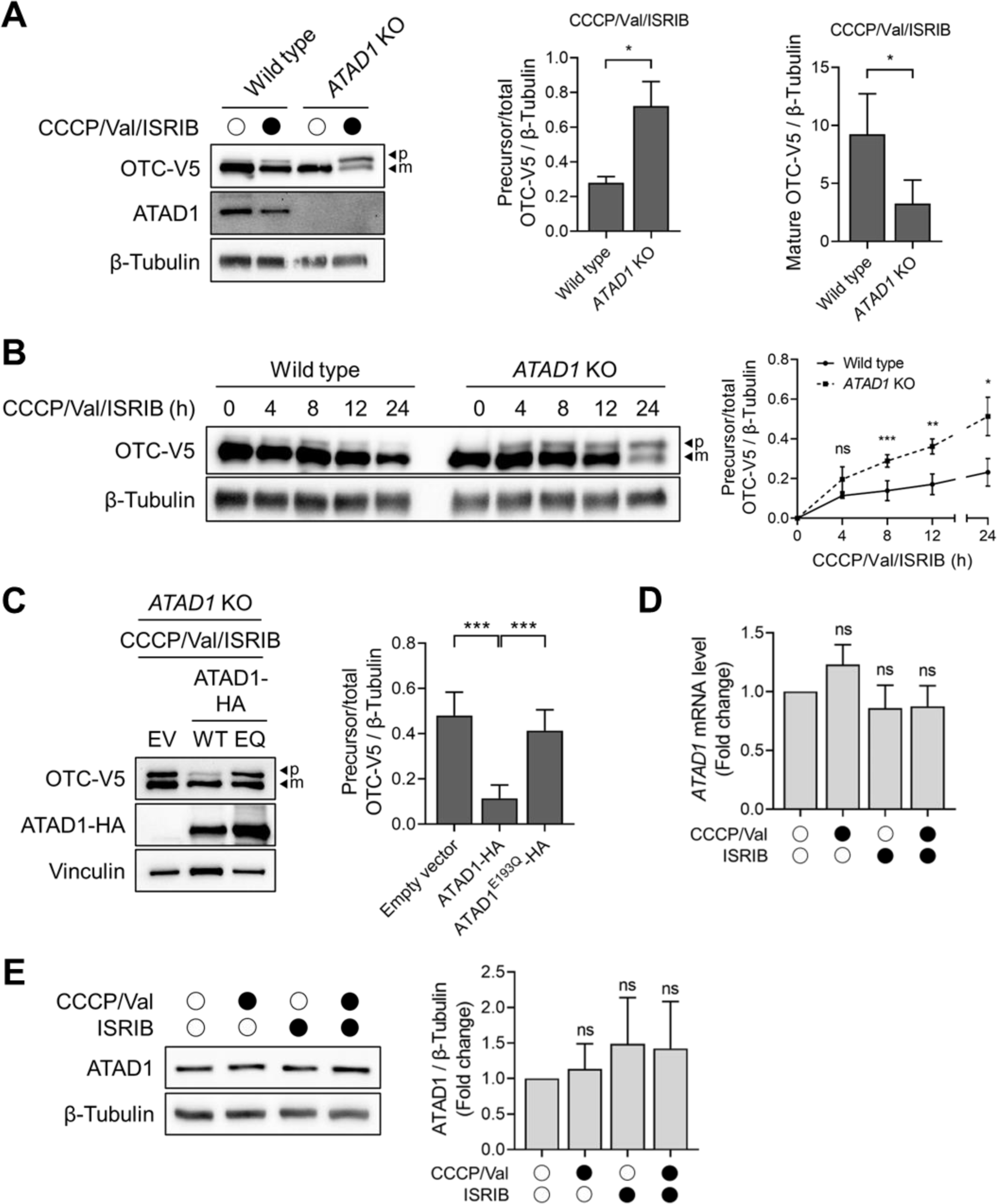
ATAD1 regulates protein quality control of mitochondrial precursors and prevents their accumulation. (**A-B**) Western blot analysis of wild type and *ATAD1* KO HEK293T cells transiently expressing OTC-V5. Mitochondrial protein import stress was induced by CCCP (10 µM), valinomycin (Val) (1 µM), and ISRIB (0.5 µM) for 24 hours (A) or for the indicated time periods (B). Quantifications of OTC-V5 precursor levels normalized to total OTC-V5, as well as mature OTC-V5, to β-tubulin are shown (*n*=3). Data are mean ± SD. Paired t-test was used (*ns* = not significant, * = *p* < 0.05, ** = *p* < 0.01, *** = *p* < 0.001). (**C**) *ATAD1* KO HEK293T cells were co-transfected with OTC-V5 and either empty vector (EV), ATAD1-HA (WT), or ATAD1^E193Q^-HA (EQ). Mitochondrial protein import stress was induced as in A. Bar graph shows densitometry of OTC-V5 precursor levels normalized to total OTC-V5 and vinculin (*n*=4). Data are mean ± SD. Paired t-test was used (*** = *p* < 0.001). (**D-E**) HEK293T cells were untreated or treated for 24 hours with CCCP/valinomycin, ISRIB, or both CCCP/valinomycin and ISRIB (same concentrations as in A). *ATAD1* mRNA level was analyzed by means of quantitative reverse transcription polymerase chain reaction (RT-PCR) and quantified by normalization to *GAPDH* (D). ATAD1 protein level was analyzed by Western blot and quantified by normalization to β-tubulin (E). Quantifications are presented as fold change, for which the untreated condition was set to 1 (*n*=3). Data are mean ± SD. Paired t-test was used (*ns* = not significant). *p*, precursor; *m*, mature protein.

To gain further insight on how ATAD1 is regulated, we examined its expression levels under different conditions. Unlike known ISR target genes (*ASNS* and *PCK2*)^16^, we detected no changes in the mRNA level of *ATAD1* when mitochondrial protein import was impaired by CCCP/valinomycin (**Figs. 3D and S3**). Similar results were obtained when ATAD1 protein levels were examined (**Fig. 3E**). A potential role for the ISR in controlling ATAD1 expression was also excluded using ISRIB alone or in combination with CCCP/valinomycin (**Figs. 3D, 3E and S3**).

### ATAD1 associates with TOM and mitochondrial precursors during protein import stress

ATAD1’s impact on mitochondrial precursor accumulation suggests a direct role for this ATPase in the clearance of TOM channels. We therefore tested whether ATAD1 associates with the TOM complex. Immunoprecipitation of ATAD1-HA from protein import stressed cells pulled down the TOM complex subunits, TOMM20 and TOMM40 (**Fig. 4A**). This association was also detected under non-stress conditions, albeit to a lesser degree, suggesting that the interaction between ATAD1 and TOM is stabilized when import channels are clogged. We next tested whether ATAD1 interacts with OTC precursor upon protein import stress. OTC-V5 was co-immunoprecipitated with ATAD1-HA in cells under protein import stress and vice versa (**Figs. 4B and 4C**). Importantly, this interaction was not detected in untreated cells.

**Figure 4.**
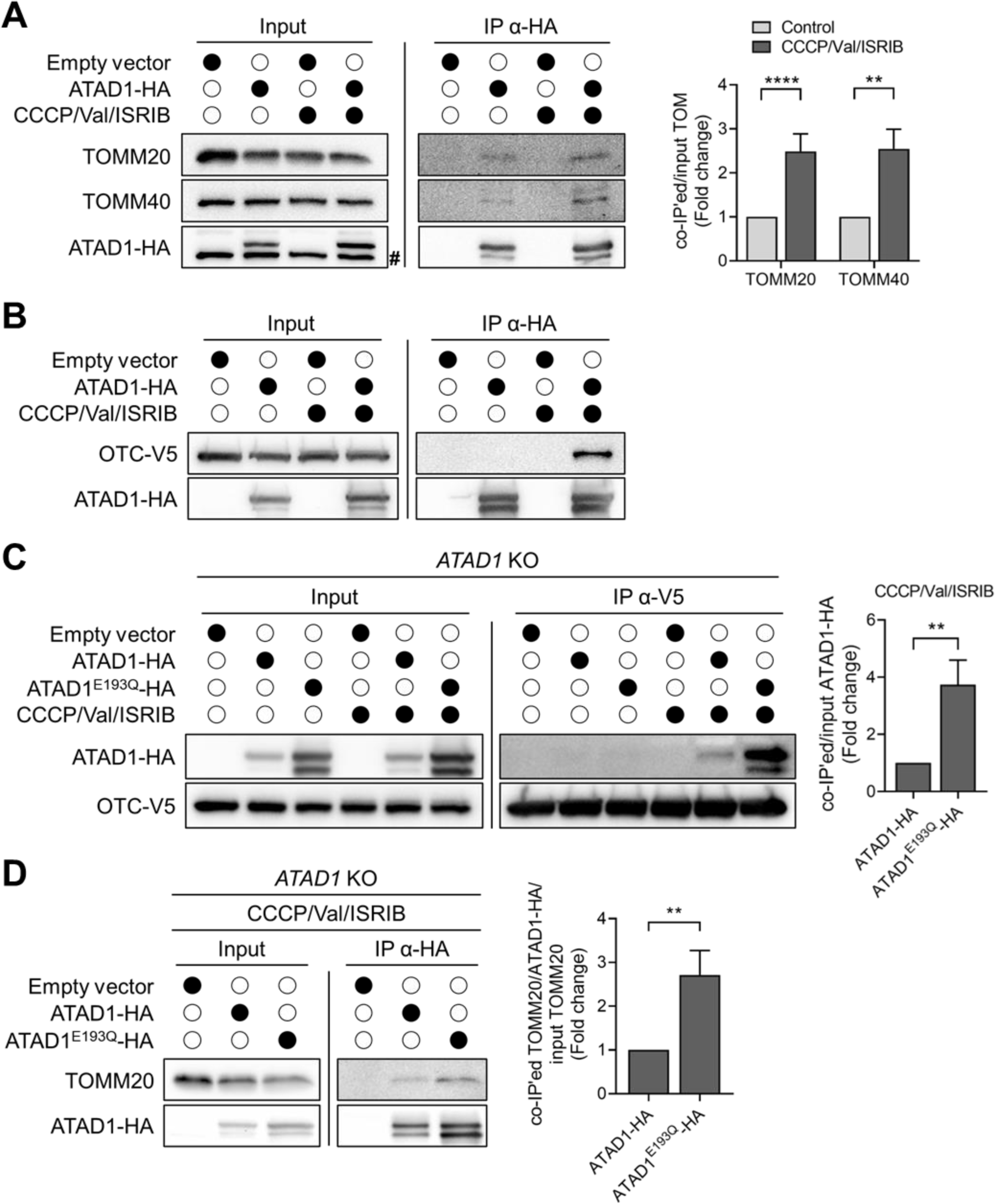
ATAD1 associates with mitochondrial precursors and TOM during protein import stress. (**A**) HEK293T cells were transfected with either empty vector or ATAD1-HA. Mitochondrial protein import stress was induced by CCCP (10 µM), valinomycin (Val) (1 µM), and ISRIB (0.5 µM) for 24 hours. ATAD1-HA was immunoprecipitated using HA antibodies. The pound symbol (#) indicates an unspecific band. Quantification of co-immunoprecipitated TOMM20 and TOMM40 normalized to their respective inputs are presented as fold change, for which the control condition was set to 1 (*n*≥4). Control conditions were normalized to 1. Data are mean ± SD. Paired t-test was used (** = *p* < 0.01, **** = *p* < 0.0001). (**B**) Same as in A. HEK293T cells stably expressing OTC-V5 were used. (**C**) OTC-V5 was immunoprecipitated from *ATAD1* KO HEK293T cells stably expressing OTC-V5, transfected with empty vector, ATAD1-HA, or ATAD1^E193Q^-HA. Mitochondrial protein import stress was induced as in A. Quantification of co-immunoprecipitated ATAD1 normalized to their respective input are presented as fold change, for which the ATAD1-HA expressing cells was set to 1 (*n*=4). Data are mean ± SD. Paired t-test was used (** = *p* < 0.01). (**D**) ATAD1-HA was immunoprecipitated from *ATAD1* KO HEK293T cells transfected with empty vector, ATAD1-HA, or ATAD1^E193Q^-HA. Mitochondrial protein import stress was induced as in A. Quantification of the IP is presented on the right. Co-immunoprecipitated TOMM20 was normalized to co-immunoprecipitated ATAD1-HA and TOMM20 input. Levels are presented as fold change, for which the ATAD1-HA expressing cells was set to 1 (*n*=4). Data are mean ± SD. Paired t-test was used (** = *p* < 0.01).

To confirm that OTC is a substrate of ATAD1, we compared the interaction of OTC with wild type versus a “substrate-trap” mutant (E193Q) of ATAD1. As ATAD1^E193Q^ is catalytically inactive, it is expected to remain bound to substates for a longer period. Indeed, immunoprecipitation of OTC-V5 revealed that it is more strongly associated with ATAD1^E193Q^-HA than with wild type ATAD1-HA (**Fig. 4C**). To eliminate the effect of endogenous ATAD1, this analysis was done in *ATAD1* knockout cells. Note that differences between ATAD1 and ATAD1^E193Q^ protein levels were consistently observed in our assays and were taken under account in the co-IP quantification (**Fig. 4C**). The inability of ATAD1^E193Q^ to hydrolyze ATP also led to a stabilized association with the TOM complex, suggesting that the interaction of ATAD1 with its substrates occurs while they are still stalled inside the translocase (**Fig. 4D**). TOMM20 protein levels were not affected by ATAD1, excluding the possibility that ATAD1 regulates the degradation of TOM (**Fig. S4**). Together, these data indicate a role for ATAD1 in extracting mitochondrial precursors that clog the outer membrane translocase.

### A cellular model with intrinsic mitochondrial defects exhibits impaired protein import

So far, our findings suggest that both ISR-mediated translational attenuation and ATAD1-mediated extraction prevent damage caused by mitochondrial protein import stress. However, as our assays were primarily conducted using harsh drug-induced treatments, it remains unclear whether these cellular surveillance mechanisms are relevant in physiological systems. To identify a cellular model with intrinsic defects in mitochondrial protein import, we surveyed various human cell lines and assessed the basal import of OTC into mitochondria. A mild accumulation of OTC-V5 precursor in HT1080 fibrosarcoma cells was observed, suggesting that mitochondrial dysfunction and inefficient protein import characterize this cell line (**Fig. 5A**). As mitochondrial damage induces the ISR, we asked whether a chronic activation of this response is a feature of HT1080 cells^16^. Indeed, a significantly higher level of phospho-eIF2α was detected in untreated HT1080 cells in comparison to other cell lines (**Fig. 5B**). Importantly, treating HT1080 cells with ISRIB for 24 hours exacerbated the accumulation of OTC-V5 precursor, supporting a role for the ISR in the quality control of un-imported mitochondrial proteins (**Fig. 5C**). Yet, inhibition of the ISR was not sufficient for detecting precursors in other cell lines. Growth rate measurements in the presence or absence of ISRIB revealed that the ISR is beneficial for the fitness of HT1080 cells (**Fig. 5D**). As expected, ISRIB treatment had no impact on the growth of HEK293T cells, which do not exhibit basal ISR activation. Altogether, these results support both a basal defect in mitochondrial protein import and a protective chronic activation of the ISR in HT1080 cells.

**Figure 5.**
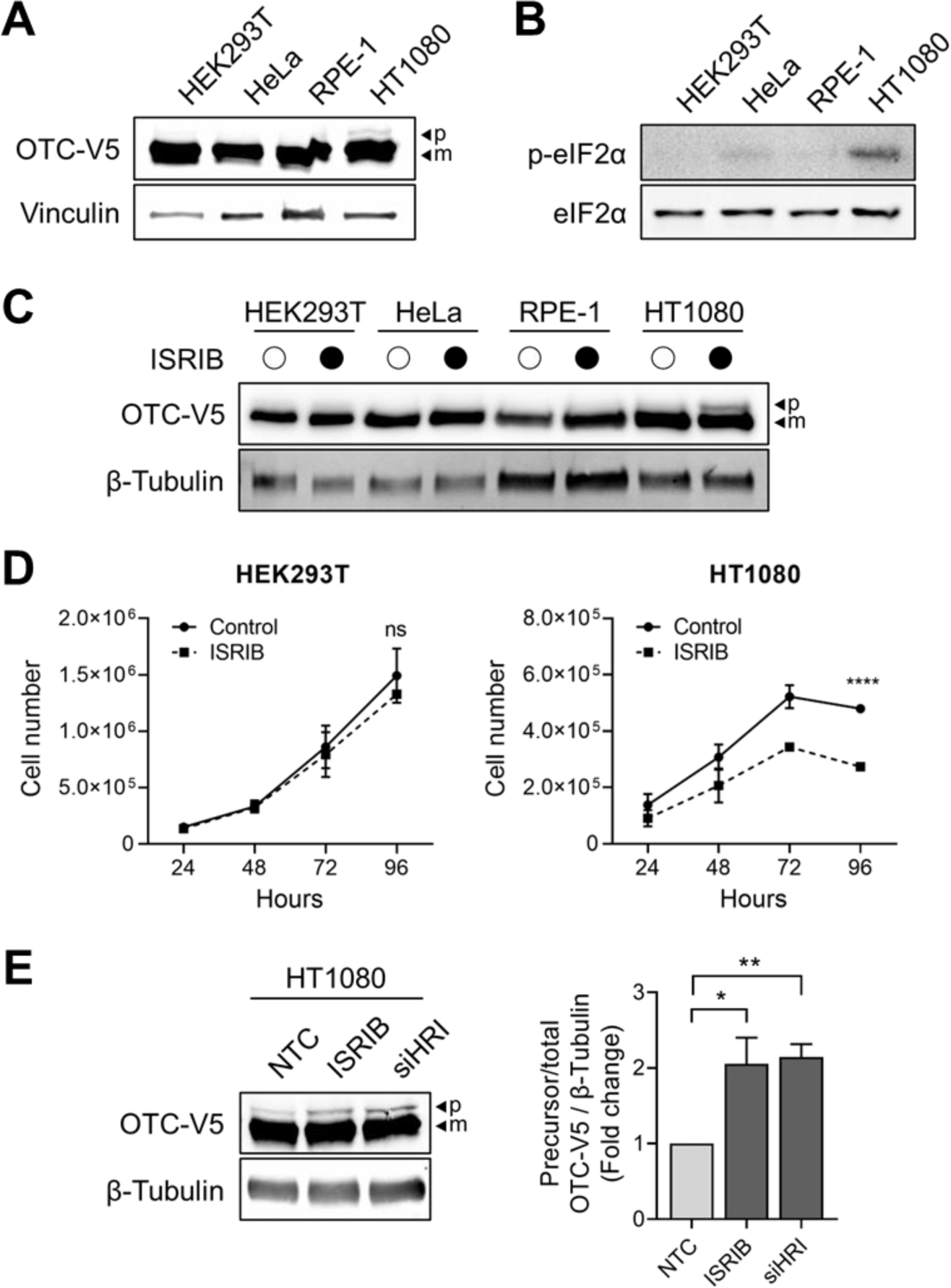
Characterization of HT1080 cell line as a cellular model with intrinsic mitochondrial protein import defects. (**A**,**C**) Western blot analysis of OTC-V5 stably expressing HEK293T, HeLa, RPE-1, and HT1080 cells untreated (A) or treated with ISRIB (0.5 µM) for 24 hours (C). (**B**) Western blot analysis of p-eIF2α level in HEK293T, HeLa, RPE-1, and HT1080 cell lines. Bar graph shows fold change in p-eIF2α level normalized to total eIF2α level, for which the HEK293T cells was set to 1 (*n*=4). Data are mean ± SD. Paired t-test was used (* = *p* < 0.05, ** = *p* < 0.01). (**D**) Growth curves of HEK293T and HT1080 cell lines untreated or treated with ISRIB (0.5 µM) for the indicated time periods. Data are mean ± SD of 3 biological replicates, consisting of 3 technical replicates each (*ns* = not significant, **** = *p* < 0.001). (**E**) OTC-V5 stably expressing HT1080 cells were treated with ISRIB (0.5 µM) for 24 hours or transfected with non-targeting control siRNA (NTC) or a pool of *HRI* siRNAs (si*HRI*) for 48 hours. Quantifications of OTC-V5 precursor levels normalized to total OTC-V5 and β-tubulin are presented as fold change, for which the NTC condition was set to 1 (*n*=3). Data are mean ± SD. Paired t-test was used (* = *p* < 0.05, ** = *p* < 0.01). *p*, precursor; *m*, mature protein.

The induction of the ISR can be triggered by various insults, including amino acid deprivation and viral infection, as well as endoplasmic reticulum or mitochondrial stress^15,16^. Different eIF2α kinases are responsible for triggering the ISR in response to different stress stimuli, with the heme-regulated inhibitor (HRI) kinase acting downstream of mitochondrial damage^15,36,37^. To verify that the ISR activation in HT1080 cells is triggered by mitochondrial stress, we silenced the expression of HRI using siRNA. Knockdown of HRI led to a significant increase in the accumulation of OTC precursor, mimicking the effect of ISRIB (**Fig. 5E**). We conclude that protein import and possibly other mitochondrial functions are disrupted in HT1080 cells and are moderated, at least partially, by the ISR.

### ATAD1 alleviates growth defects in cells with impaired mitochondrial protein import

Identifying the HT1080 cell line as a cellular model with defective protein import enabled the investigation of ATAD1’s contribution to cellular health. Protein quantification analysis revealed low abundance of ATAD1 in HT1080 cells compared to other cell lines (**Fig. 6A**). We therefore asked whether increasing the levels of ATAD1 by lentiviral transduction (ATAD1^↑^) could impact the accumulation of mitochondrial precursors. Assessment of OTC-V5 levels revealed that while ATAD1^↑^ expression was insufficient to restore mitochondrial protein import in HT1080 cells, it led to a 45% reduction in the accumulation of OTC-V5 precursor (**Fig. 6B**). Importantly, cellular fitness was positively affected by increased ATAD1 levels, as evidenced by the improved proliferation of ATAD1^↑^ stably expressing cells, in comparison to parental HT1080 cells (**Fig. 6C**). In contrast, ATAD1^E193Q↑^ stable expression had no impact on proliferation, presumably since the endogenous ATAD1 was still present in these cells. ATAD1^↑^ expression had no effect on HEK293T cells that do not exhibit a mitochondrial chronic stress. Together, these data indicate a central role for ATAD1 in mitochondrial protein import surveillance and cellular health.

**Figure 6.**
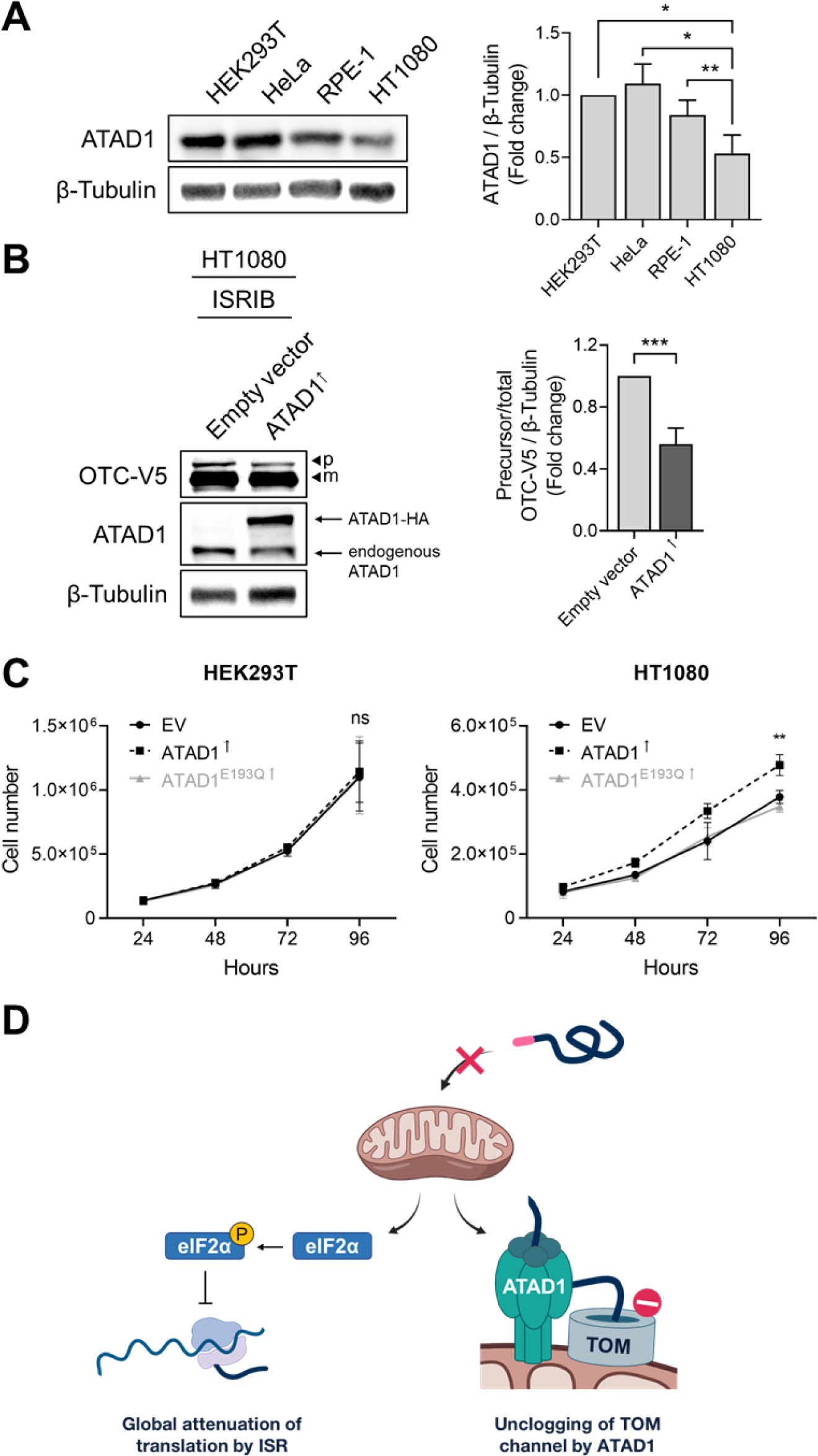
ATAD1 impacts proliferation rate of cells with impaired mitochondrial protein import. (**A**) Western blot analysis of endogenous ATAD1 protein from HEK293T, HeLa, RPE-1, and HT1080 cells. Bar graph shows fold change in ATAD1 level normalized to β-tubulin, for which the HEK293T cells was set to 1 (*n*=3). Data are mean ± SD. Paired t-test was used (* = *p* < 0.05, ** = *p* < 0.01). (**B**) Western blot analysis of empty vector or ATAD1-HA (ATAD1^↑^) stably expressing HT1080 cells transiently transfected with OTC-V5 and treated with ISRIB for 24 hours. *p*, precursor; *m*, mature protein. Quantifications of OTC-V5 precursor levels normalized to total OTC-V5 and β-tubulin are presented as fold change, for which the empty vector condition was set to 1 (*n*=6). Data are mean ± SD. Paired t-test was used (*** = *p* < 0.001). (**C**) Growth curves of HEK293T and HT1080 cell lines stably expressing empty vector, ATAD1^↑^, or ATAD1^E193Q↑^. Data are mean ± SD of 3 biological replicates, consisting of 2 technical replicates each (*ns* = not significant, ** = *p* < 0.01). (**D**) Overall schematic of proposed model. The stalling of un-imported proteins inside TOM can be alleviated by attenuating global protein translation via ISR, as well as removal of un-imported proteins from TOM by ATAD1.

## Discussion

Here, we investigate the consequences and cellular response to defects in mitochondrial protein import in mammalian cells. We demonstrate that when import is impaired, mitochondrial proteins can accumulate outside the organelle and as translocation intermediates inside TOM. Under these conditions, activation of the ISR alleviates the accumulation of un-imported mitochondrial precursors, including those stalled inside TOM, by attenuating global protein translation. Moreover, we show that the ATPase, ATAD1, has a role in extracting stalled precursors from the mitochondrial translocase. We propose a model where different mechanisms help cells cope with damage caused by impaired mitochondrial protein import. As the first line of defense, the ISR-mediated translation inhibition prevents proteotoxicity and the overwhelming of cellular protein quality control pathways. More specific mechanisms, such as ATAD1-mediated unclogging of the entry gate into the mitochondria, act in parallel. We suggest that by clearing TOM, ATAD1 supports some level of protein import while the damage persists and aids in rapid recovery after the damage subsides (**Fig. 6D**). This model is consistent with previous work from budding yeast, where 1) mitochondrial protein import defects cause inhibition of global translation^17,18^, and 2) Msp1 (ATAD1 homolog) unclogs TOM^21,39^. Some evidence for TOM clogging also exists in mammalian cells, however these are limited to synthetically designed cloggers, mistargeted APP, or pathogenic mutants of mitochondrial proteins^26,27,40^. Our work is the first to demonstrate the ability of native mitochondrial proteins to clog the human translocase. This finding further strengthens the relevance of mitochondrial clogging to physiological and disease conditions with damaged mitochondria.

ATAD1’s function is not restricted to TOM clearance but also includes the removal of mistargeted ER or peroxisomal proteins inserted into the mitochondrial outer membrane^28–30^. How the different functions of ATAD1 are regulated is yet to be revealed. As the expression of ATAD1 is not altered by mitochondrial protein import stress or the ISR, it is likely that its TOM-unclogging activity is regulated on the mitochondrial surface. Our data indicate a stronger association of ATAD1 with the TOM complex during protein import stress, suggesting a possible recruitment of ATAD1 to TOM when protein import is inhibited. However, we cannot exclude the possibility that the association with TOM is a result of ATAD1’s interaction with stalled substrates rather than direct binding to TOM subunits. Further studies are required to answer these questions as well as to identify ATAD1 substrates and investigate how they are recognized. Other factors, such as the AAA Cdc48/p97, are also known to be involved in the extraction of stalled precursors in yeast^24^. Although a similar role for these factors in mammals is yet to be defined, it is possible that they act in parallel to ATAD1 and extract other subsets of substrates.

The fate of stalled precursors following ATAD1-mediated extraction is still unclear. Previous reports, focused mainly on extraction of mistargeted ER/peroxisomal proteins by Msp1, suggested both redirection to the proper organelle and degradation as possible outcomes^24,28–30,39,41^. As protein degradation and translation require energy, it is plausible that degradation of substrates is used as a last resort. Therefore, ATAD1-mediated unfolding and TOM extraction could provide stalled precursors a second chance for translocation^42^. Our finding that proteasomal inhibition does not lead to further accumulation of OTC precursor might support the re-entry model over degradation. An additional support for this model comes from our observation that mature OTC levels were decreased in *ATAD1* KO cells. Regardless of the model, ATAD1 activity can help protect the cytosol from proteotoxicity by preventing accumulation of un-imported proteins.

We describe a cellular model that exhibits chronic mitochondrial stress (HT1080 cell line) to demonstrate the beneficial impact of ISR on cell fitness. Two outcomes of ISR activation are global protein translation attenuation and induction of downstream transcriptional stress programs^15^. It remains unclear what the contribution is for each of these branches on the accumulation of mitochondrial precursors and cellular health. ISR-mediated stress responses, which depend on transcription factors such as ATF4 and ATF5, were demonstrated to affect mitochondrial health and are likely to have a major role in HT1080 cell homeostasis^16,43^. The nature of the mitochondrial defect that triggers the ISR in the fibrosarcoma HT1080 cell model is unknown. It is possible that this defect is characteristic of certain types of cancers. Thus, combating impaired mitochondrial protein import might be an essential mechanism for the viability of these cancer cells. It is intriguing to reveal how the ISR activation is regulated in these cancers and under other conditions with chronic defects in mitochondrial protein import. Fluctuation in ISR activation might occur to balance translation of housekeeping proteins with prevention of proteotoxicity caused by un-imported proteins. Other disease conditions, as well as physiological conditions, are associated directly or indirectly with a slowdown in mitochondrial protein import^6,8,10,19,23,44^. For example, during development, the import of *ATF5* and its worm homolog *ATFS-1* serves as a reporter for mitochondrial capacity and regulates the expansion of the mitochondrial network^23,43^. Our finding suggests a potential role for both ISR and ATAD1 in recovering from such conditions.

## Materials and methods

### Cell culture procedures and treatment

All cells were grown in an incubator at 5% CO_2_ and 37°C. HEK293T and HEK293FT cells were cultured in Dulbecco’s modified Eagle’s medium (DMEM) (Gibco), and HT1080 cells were cultured in Roswell Park Memorial Institute (RPMI) 1640 (Gibco). Cell media was supplemented with 10% Fetal Bovine Serum (Gibco), 2 mM L-glutamine (Gibco), and 100 U/mL penicillin-streptomycin (Gibco).

Cell transfections were done using X-tremeGENE HP DNA Transfection Reagent (Roche). Cells were harvested 48 hours following transfection. CCCP (Fisher Scientific), valinomycin (Invitrogen), and ISRIB (Sigma-Aldrich) were administered at 10 µM, 1 µM, and 0.5 µM, respectively, for 24 hours. For siRNA knockdown experiments, cells were transiently transfected with 25 nM ON-TARGETplus human *HRI* (*EIF2AK1*) siRNA SMARTpool (Dharmacon) or ON-TARGETplus Non-targeting (control) pool (Dharmacon) using DharmaFECT 4 transfection reagent (Dharmacon) according to the manufacturer’s protocol.

Stable expression of OTC and ATAD1 in HEK293T/HT1080 cells was obtained using lentiviral transduction. Lentiviral production was done by co-transfecting HEK293FT cells with 3^rd^ generation lentiviral packaging system and the plasmid of interest (pRSV-Rev, pMDLg/pRRE, pMD2.G, and pLV-EF1a-IRES-Hygro, in 1:2:1.2:3.4 molar ratio) and changing to fresh media ∼14 hours post-transfection. Supernatant of HEK293FT cells containing fresh lentivirus was collected 48 hours post-transfection. For infection, HEK293T or HT1080 cells were grown with 10 mL filtered viral supernatant containing 1 µg/mL polybrene (EMD Millipore). HEK293T and HT1080 cells were incubated for 24 hours before the addition of 400 µg/mL and 100 µg/mL hygromycin (Roche) for selection, respectively.

### Plasmids

To construct the OTC-V5 expression plasmid, rat *OTC* (from Addgene plasmid #71877) was cloned into the lentiviral backbone pLV-EF1a-IRES-Hygro (Addgene plasmid #85134). To construct the ATAD1-HA plasmid, human *ATAD1* was amplified from HEK293T cDNA and cloned into the pKH3 vector (Addgene plasmid #12555). The ATAD1 Walker B mutation (ATAD1^E193Q^) was introduced by generating gene blocks (gBlocks) and cloning into the PstI and Bsu36I sites of the pKH3-ATAD1-HA vector. The PISD-FLAG plasmid was generated by amplifying human *PISD* from cDNA and cloning it into the BglII and SalI sites of the pmCherry-N1 vector (Clontech). The C-terminal mCherry tag was subsequently replaced with 3xFLAG tag using the AgeI and NotI restriction sites. The COQ7-HA plasmid was constructed by PCR amplification of human *COQ7* from cDNA and cloning into the pKH3 vector. All cloning was performed using Gibson assembly (NEB). The ATP5G1-HA expression plasmid was a gift from Dr. Ramanujan S. Hegde (MRC Laboratory of Molecular Biology, Cambridge, UK). Lentiviral packaging system: pRSV-Rev (Addgene plasmid #12253), pMDLg/pRRE (Addgene plasmid #12251), and pMD2.G (Addgene plasmid #12259).

### Generation of KO cell lines by CRISPR/Cas9

Two gRNAs targeting the first exon of human *ATAD1* (5’-CCGACTCAAAGGACGAGAAA-3’ and 5’-TGACATACTTTACTATCAAA-3’), designed using the online tool CHOPCHOP^45^, were cloned into the pU6-sgRNA-CBh-Cas9-PGK-Venus vector. Forty-eight hours following transfection of the vector into HEK293T, single Venus expressing cells were sorted by flow cytometry into 96-well plates. Gene knockout was validated by Western blot using a monoclonal ATAD1 antibody (BioLegend), as well as PCR amplification of the genomic region flanking the targeting sequence.

### Protein analysis and antibodies

Cells were lysed using RIPA buffer (Thermo Fisher Scientific) supplemented with cOmplete protease inhibitors (Roche). PhosSTOP phosphatase inhibitor (Roche) was added for p-eIF2α analysis. The following primary antibodies were used: mouse anti-ATAD1 (1:500, BioLegend), rabbit anti-β-Tubulin (1:5000, Absolute Antibody), rabbit anti-COXIV (1:1000, Cell Signaling Technology), mouse anti-eIF2α (1:1000, Invitrogen), rabbit anti-p-eIF2α (1:500, Cell Signaling Technology), mouse anti-HA (1:1000, BioLegend), rabbit anti-HA (1:1000, Abcam), rabbit anti-TOMM20 (1:1000, Abcam), rabbit anti-TOMM40 (1:1000, Abcam), mouse anti-V5 (1:1000, Abcam), rat anti-V5 (1:1000, Abcam), and mouse anti-Vinculin (1:1000, Sigma-Aldrich). Secondary antibodies: sheep anti-mouse HRP-conjugated (1:10,000, Thermo Fisher Scientific), donkey anti-rabbit HRP-conjugated (1:10,000, Thermo Fisher Scientific), donkey anti-rat HRP-conjugated (1:10,000, Jackson ImmunoResearch), goat anti-mouse DyLight 800 (1:15,000, Invitrogen), and goat anti-rabbit DyLight 680 (1:15,000, Invitrogen).

### Mitochondrial isolation and protease protection assay

Cells were resuspended in cold homogenization buffer (250 mM sucrose, 250 mM HEPES/KOH pH 7.4, 1 mM EDTA) supplemented with cOmplete protease inhibitors (Roche) and lysed by Dounce homogenization on ice. Mitochondria were separated from cytosol by centrifugation of post-nuclear fraction at 4°C (10,000 x g for 10 mins). Mitochondrial fractions were subjected to various concentrations (0.25-1 mg/mL) of proteinase K (Sigma-Aldrich) for 30 mins at 37°C. The reaction was stopped (15 mins on ice) using 4 mM phenylmethylsulfonyl fluoride (PMSF; Thermo Fisher Scientific).

### Coimmunoprecipitation

Cells were lysed in 50 mM Tris pH 7.5, 250 mM NaCl, 1 mM EDTA, and 5% IGEPAL CA-630 supplemented with cOmplete protease inhibitors (Roche), and incubated on ice for 15 mins. Following clearance (16,000 x g for 10 min at 4°C), the lysates were incubated with antibody-conjugated Protein G magnetic beads (Invitrogen) for 2.5 hours at 4°C. The beads were then washed 5x in 200 µL cold wash buffer (1X PBS and 0.1% Tween-20), and bound protein was eluted by addition of SDS sample buffer and boiling.

### RNA isolation and quantitative PCR

Total RNA was isolated from cells using the RNAspin Mini kit (Cytiva) following the manufacturer’s instructions. RNA (750 ng) was used to generate cDNA using the SuperScript III First-Strand Synthesis System (Invitrogen). Quantitative PCR was performed using the PerfeCTa SYBR Green FastMix (Quantabio) according to the manufacturer’s instructions. All data were normalized to *GAPDH* transcript levels and are presented as fold increase of control conditions. The following primers were used in this study: *ATAD1* forward 5’–TTTCTGCCCCTGGGTTAACAT–3’ and reverse 5’–TCTGCCTGTTTCTGAGCTTCTA–3’; *ASNS* forward 5’– ATCACTGTCGGGATGTACCC–3’ and reverse 5’–TGATAAAAGGCAGCCAATCC–3’; *PCK2* forward 5’–CATCCGAAAGCTCCCCAAGT–3’ and reverse 5’–GCTCTCTACTCGTGCCACAT–3’; and *GAPDH* forward 5’–GTCTCCTCTGACTTCAACAGCG–3’ and reverse 5’– ACCACCCTGTTGCTGTAGCCAA–3’.

### Quantification and statistical analysis

Band intensities on Western blots were quantified using Image Lab (Bio-Rad Laboratories). All statistical data were calculated and graphed using GraphPad Prism 8. Statistical significance was assessed using two-tailed paired Student’s *t*-test. All reported *n* numbers refer to biological replicates. A P value of <0.05 was considered statistically significant (\**p* < 0.05, \*\**p* < 0.01, \*\*\**p* < 0.001 and \*\*\*\**p* < 0.0001). All error bars are expressed as mean ± SD.

## Acknowledgments

The work was supported by the Natural Sciences and Engineering Research Council of Canada (grant RGPIN-2020-05204 to H.W.), the Canadian Institutes of Health Research (grant PJT-180426 to H.W.), and Michael Smith Health Research BC (SCH-2021-1524 to H.W.). J.K. was supported by fellowships from the Canadian Institutes of Health Research and Korean Canadian Scholarship Foundation.

## Author contributions

J.K. and H.W. designed and interpreted the experiments. J.K., M.G., H.W. drafted the manuscript. J.K., M.G., L.Z. acquired and analyzed the data.

---

**Figure S1.**
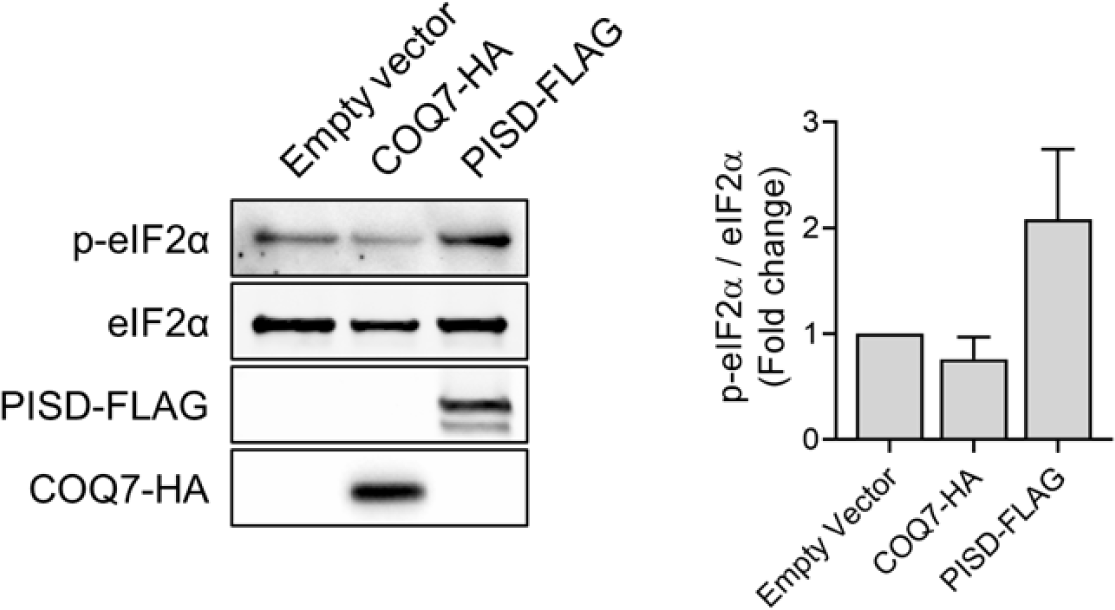
Overexpression of PISD activates the ISR. Western blot analysis of HEK293T cells transiently transfected with either empty vector, COQ7-HA (control), or PISD-FLAG. Bar graph shows fold change in p-eIF2α level normalized to total eIF2α level, for which the empty vector condition was set to 1 (*n*=2).

**Figure S2.**
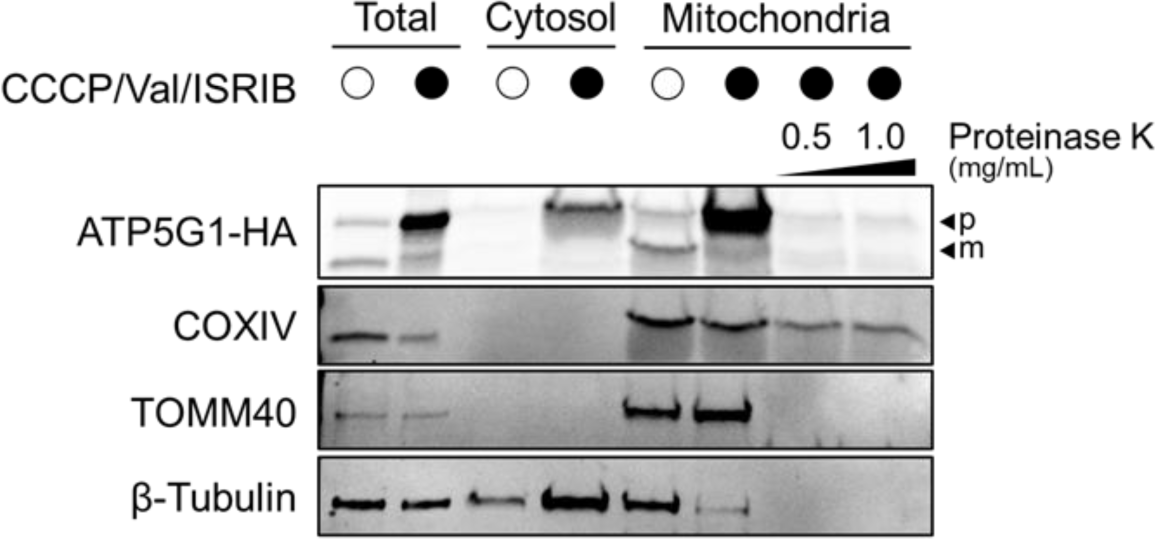
ATP5G1-HA precursors accumulate in both the cytosol and mitochondria upon protein import stress. Mitochondrial and cytosolic fractions were enriched by differential centrifugation from ATP5G1-HA transiently expressing HEK293T cells untreated or treated with CCCP (10 µM), valinomycin (Val) (1 µM), and ISRIB (0.5 µM) for 24 hours to induce protein import stress. Mitochondrial fractions were treated with 0.5 mg/mL or 1.0 mg/mL of proteinase K. TOMM40 was used as a mitochondrial outer membrane marker. COXIV was used as a mitochondrial matrix marker.

**Figure S3.**
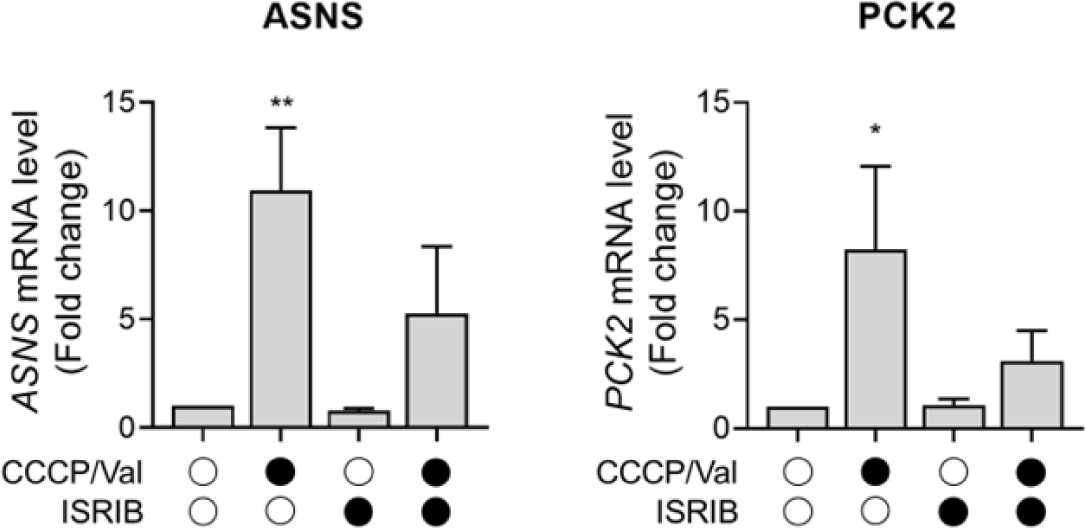
Upregulation of ISR target genes upon defects in mitochondrial protein import. HEK293T cells were untreated or treated for 24 hours with CCCP/valinomycin, ISRIB, or both CCCP/valinomycin and ISRIB (CCCP 10 µM, valinomycin (Val) 1 µM, ISRIB 0.5 µM). *ASNS* and *PCK2* mRNA levels were analyzed by means of quantitative reverse transcription polymerase chain reaction (RT-PCR) and quantified by normalization to *GAPDH*. Quantifications are presented as fold change, for which the untreated condition was set to 1 (*n*=4). Data are mean ± SD. Paired t-test was used (* = *p* < 0.05, ** = *p* < 0.01).

**Figure S4.**
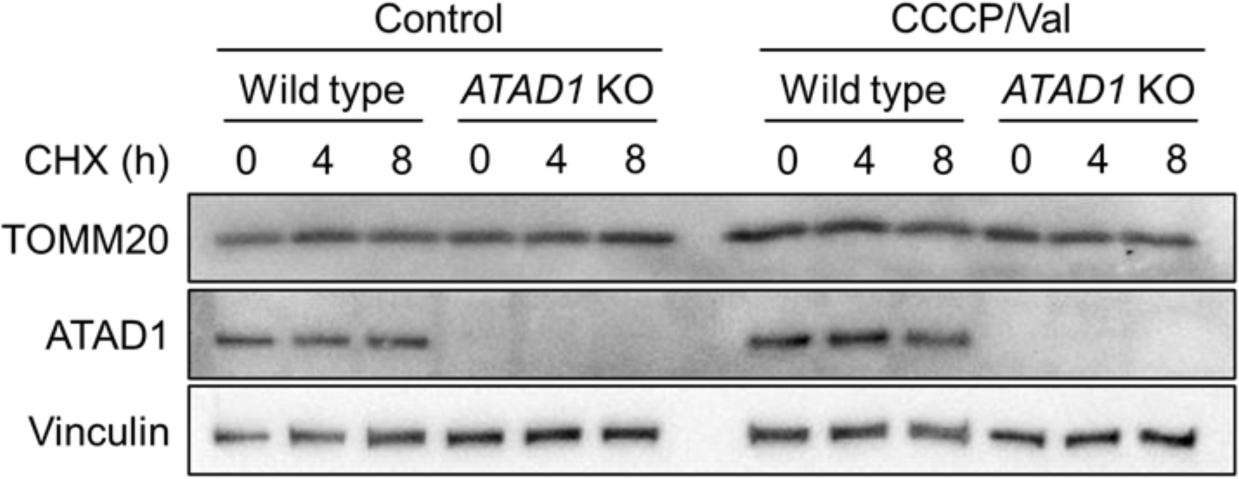
ATAD1 activity does not impact the half-life of TOM. Western blot analysis of TOMM20 protein level in wild type and *ATAD1* KO HEK293T cells. Cells were untreated (control) or treated with CCCP (10 µM) and valinomycin (1 µM) (CCCP/Val). Cycloheximide (CHX) (0.25 mg/mL) chase was performed to assess protein stability following 24 hours of CCCP/valinomycin treatment.

## References

1. Nunnari, J. & Suomalainen, A. Mitochondria: In sickness and in health. Cell 148, 1145– 1159 (2012).

2. Rath, S. et al. MitoCarta3.0: An updated mitochondrial proteome now with sub-organelle localization and pathway annotations. Nucleic Acids Res. 49, D1541–D1547 (2021).

3. Morgenstern, M. et al. Quantitative high-confidence human mitochondrial proteome and its dynamics in cellular context. Cell Metab. 33, 2464–2483.e18 (2021).

4. Busch, J. D., Fielden, L. F., Pfanner, N. & Wiedemann, N. Mitochondrial protein transport: Versatility of translocases and mechanisms. Mol. Cell 83, 890–910 (2023).

5. Frazier, A. E., Thorburn, D. R. & Compton, A. G. Mitochondrial energy generation disorders: Genes, mechanisms, and clues to pathology. J. Biol. Chem. 294, 5386–5395 (2019).

6. MacKenzie, J. A. & Payne, R. M. Mitochondrial protein import and human health and disease. Biochim. Biophys. Acta - Mol. Basis Dis. 1772, 509–523 (2007).

7. Nicolas, E., Tricarico, R., Savage, M., Golemis, E. A. & Hall, M. J. Disease-Associated Genetic Variation in Human Mitochondrial Protein Import. Am. J. Hum. Genet. 104, 784– 801 (2019).

8. Palmer, C. S., Anderson, A. J. & Stojanovski, D. Mitochondrial protein import dysfunction: mitochondrial disease, neurodegenerative disease and cancer. FEBS Lett. 595, 1107– 1131 (2021).

9. Boos, F., Labbadia, J. & Herrmann, J. M. How the Mitoprotein-Induced Stress Response Safeguards the Cytosol: A Unified View. Trends Cell Biol. 30, 241–254 (2020).

10. Nowicka, U. et al. Cytosolic aggregation of mitochondrial proteins disrupts cellular homeostasis by stimulating the aggregation of other proteins. Elife 10, 1–27 (2021).

11. Song, J., Herrmann, J. M. & Becker, T. Quality control of the mitochondrial proteome. Nat. Rev. Mol. Cell Biol. 22, 54–70 (2021).

12. Ng, M. Y. W., Wai, T. & Simonsen, A. Quality control of the mitochondrion. Dev. Cell 56, 881–905 (2021).

13. Shpilka, T. & Haynes, C. M. The mitochondrial UPR: Mechanisms, physiological functions and implications in ageing. Nat. Rev. Mol. Cell Biol. 19, 109–120 (2018).

14. Pakos-Zebrucka, K., et al. The integrated stress response. EMBO Rep. 17, 1374–1395 (2016).

15. Costa-Mattioli, M. & Walter, P. The integrated stress response: From mechanism to disease. Science 368, (2020).

16. Quirós, P. M. et al. Multi-omics analysis identifies ATF4 as a key regulator of the mitochondrial stress response in mammals. J. Cell Biol. 216, 2027–2045 (2017).

17. Wang, X. & Chen, X. J. A cytosolic network suppressing mitochondria-mediated proteostatic stress and cell death. Nature 524, 481–484 (2015).

18. Wrobel, L. et al. Mistargeted mitochondrial proteins activate a proteostatic response in the cytosol. Nature 524, 485–488 (2015).

19. Krämer, L. et al. MitoStores: chaperone-controlled protein granules store mitochondrial precursors in the cytosol. EMBO J. 1–18 (2023) doi:10.15252/embj.2022112309.

20. Boos, F. et al. Mitochondrial protein-induced stress triggers a global adaptive transcriptional programme. Nat. Cell Biol. 21, 442–451 (2019).

21. Weidberg, H. & Amon, A. MitoCPR—A surveillance pathway that protects mitochondria in response to protein import stress. Science (80-.). 360, (2018).

22. Sutandy, F. X. R., Gößner, I., Tascher, G. & Münch, C. A cytosolic surveillance mechanism activates the mitochondrial UPR. Nature (2023) doi:10.1038/s41586-023-06142-0.

23. Shpilka, T. et al. UPRmt scales mitochondrial network expansion with protein synthesis via mitochondrial import in Caenorhabditis elegans. Nat. Commun. 12, (2021).

24. Mårtensson, C. U. et al. Mitochondrial protein translocation-associated degradation. Nature 569, 679–683 (2019).

25. Schulte, U. et al. Mitochondrial complexome reveals quality-control pathways of protein import. Nature (2023) doi:10.1038/s41586-022-05641-w.

26. Coyne, L. P. et al. Mitochondrial protein import clogging as a mechanism of disease. Elife 12, 1–42 (2023).

27. Devi, L., Prabhu, B. M., Galati, D. F., Avadhani, N. G. & Anandatheerthavarada, H. K. Accumulation of amyloid precursor protein in the mitochondrial import channels of human Alzheimer’s disease brain is associated with mitochondrial dysfunction. J. Neurosci. 26, 9057–9068 (2006).

28. Chen, Y. et al. Msp1/ATAD1 maintains mitochondrial function by facilitating the degradation of mislocalized tail-anchored proteins. EMBO J. 33, 1548–1564 (2014).

29. Okreglak, V. & Walter, P. The conserved AAA-ATPase Msp1 confers organelle specificity to tail-anchored proteins. Proc. Natl. Acad. Sci. U. S. A. 111, 8019–8024 (2014).

30. Wang, L. & Walter, P. Msp1/ATAD1 in Protein Quality Control and Regulation of Synaptic Activities. Annu. Rev. Cell Dev. Biol. 36, 141–164 (2020).

31. Itakura, E. et al. Ubiquilins Chaperone and Triage Mitochondrial Membrane Proteins for Degradation. Mol. Cell 63, 21–33 (2016).

32. Zhao, Q. et al. A mitochondrial specific stress response in mammalian cells. EMBO J. 21, 4411–4419 (2002).

33. Shakya, V. P. S. et al. A nuclear-based quality control pathway for non-imported mitochondrial proteins. Elife 10, 1–21 (2021).

34. Fu, Y., Kanshin, E., Ueberheide, B. & Sfeir, A. Mitochondrial DNA Breaks Activate an Integrated Stress Response to Reestablish Homeostasis. bioRxiv 2022.10.01.510431 (2023).

35. Schäfer, J. A., Bozkurt, S., Michaelis, J. B., Klann, K. & Münch, C. Global mitochondrial protein import proteomics reveal distinct regulation by translation and translocation machinery. Mol. Cell 82, 435–446.e7 (2022).

36. Guo, X. et al. Mitochondrial stress is relayed to the cytosol by an OMA1–DELE1–HRI pathway. Nature 579, 427–432 (2020).

37. Fessler, E. et al. A pathway coordinated by DELE1 relays mitochondrial stress to the cytosol. Nature 579, 433–437 (2020).

38. Samluk, L. et al. Cytosolic translational responses differ under conditions of severe short-term and long-term mitochondrial stress. Mol. Biol. Cell 30, 1864–1877 (2019).

39. Basch, M. et al. Msp1 cooperates with the proteasome for extraction of arrested mitochondrial import intermediates. Mol. Biol. Cell 31, 753–767 (2020).

40. Krakowczyk, M. et al. OMA1 protease eliminates arrested protein import intermediates upon depolarization of the inner mitochondrial membrane. bioRxiv 2023.06.08.543713 (2023) 10.1101/2023.06.08.543713.

41. Matsumoto, S. et al. Msp1 Clears Mistargeted Proteins by Facilitating Their Transfer from Mitochondria to the ER. Mol. Cell 76, 191–205.e10 (2019).

42. Castanzo, D. T., LaFrance, B. & Martin, A. The AAA+ ATPase Msp1 is a processive protein translocase with robust unfoldase activity. Proc. Natl. Acad. Sci. U. S. A. 117, 14970–14977 (2020).

43. Fiorese, C. J. et al. The Transcription Factor ATF5 Mediates a Mammalian Mitochondrial UPR. Curr. Biol. 26, 2037–2043 (2016).

44. Peng, M. et al. Inhibiting cytosolic translation and autophagy improves health in mitochondrial disease. Hum. Mol. Genet. 24, 4829–4847 (2015).

45. Montague, T. G., Cruz, J. M., Gagnon, J. A., Church, G. M. & Valen, E. CHOPCHOP: A CRISPR/Cas9 and TALEN web tool for genome editing. Nucleic Acids Res. 42, 401–407 (2014).

